# Improved prediction of virus-human protein-protein interactions by incorporating network topology and viral molecular mimicry

**DOI:** 10.64898/2026.02.28.708776

**Authors:** Zhiyuan Zhang, Yang Feng, Xiangxian Meng, Yousong Peng

## Abstract

The protein-protein interactions (PPIs) between viruses and human play crucial roles in viral infections. Although numerous computational approaches have been proposed for predicting virus-human PPIs, their performances remain suboptimal and may be overestimated due to the lack of benchmark dataset. To address these limitations, we first constructed a carefully curated benchmark dataset, ensuring non-overlapped PPIs and minimum sequences similarity of both human and viral proteins in the training and test sets. Based on this dataset, we developed vhPPIpred, a machine learning-based prediction method that not only incorporated sequence embedding and evolutionary information but also leveraged network topology and viral molecular mimicry of human PPIs. Comparative experiments demonstrated that vhPPIpred outperformed six state-of-the-art methods on both our benchmark dataset and three independent datasets. Ablation analysis showed that the sequence embedding and network topology features contributed most to vhPPIpred. Besides, vhPPIpred also achieved high computational efficiency, requiring relatively low runtime and memory. Finally, vhPPIpred was demonstrated to have great potential in identifying human virus receptors. In summary, this study provides a valuable benchmark dataset and an effective tool for virus-human PPI prediction, with potential applications in antiviral drug discovery and host-pathogen interaction research.

## Background

Viral infections seriously threaten human health. According to the World Health Organization (WHO), the pandemic caused by Severe Acute Respiratory Syndrome coronavirus 2 (SARS-CoV-2) has resulted in more than 777 million human infections and 7 million deaths as of February of 2025[1]. Vaccines and drugs are the primary strategies for prevention and control of viral infections. For example, palivizumab targets the fusion (F) protein of respiratory syncytial virus (RSV) to prevent human from viral infections[2], while maraviroc treats human immunodeficiency virus (HIV) by targeting the human CCR5, a receptor for HIV[3]. Investigations of protein-protein interactions (PPIs) between viruses and human will elucidate the mechanisms of viral infections and accelerate the development of antiviral strategies.

Experimental techniques for identifying PPIs can be classified into two categories: low-throughput and high-throughput approaches. Low-throughput approaches, such as pulldown assays[4], electron microscopy[5] and isothermal titration calorimetry[6], are capable of detecting a limited number of interactions with high reliability. In contrast, high-throughput approaches like yeast two-hybrid (Y2H)[7], affinity purification coupled with mass spectrometry (AP-MS)[8], and protein microarrays[9] have enabled the large-scale identification of PPIs. For instance, Calderwood et al. used Y2H to identify PPIs between Epstein-Barr virus (EBV) and human[10], while Shah et al. combined AP-MS with RNAi screening to detect dengue virus (DENV)-human PPIs[11]. These efforts have fueled the development of specialized databases of virus-human PPIs such as Viruses.STRING[12], HVIDB[13], and HVPPI[14]. However, such experimental methods remain constrained by inherent limitations including substantial time investment, high costs, labor intensity, and frequent false-positive results. Moreover, when investigating virus-host interactions, these experimental techniques require stringent biosafety conditions due to the potential risks associated with viral pathogens. These constraints hinder the application of experimental approaches in large-scale identification of virus-human PPIs.

Computational methods for predicting virus-human PPIs have become an indispensable complement to experimental approaches. They can be broadly categorized into four types: interolog mapping, domain-domain/motif interaction-based inference, structure-based method, and machine learning-based method as summarized in Lian’s review[15]. The main idea of interolog mapping is to infer unknown PPIs from known homologous PPIs[16], while its applicability is constrained by the generally low homology of viral proteins. The domain-domain/motif interaction-based inference relies on the detection of the domain-domain/motif interacting pairs in the query protein pair to infer the potential interactions[17,18], while this approach is limited by the known domain-domain/motif interaction pairs. The structure-based method includes two types: (1) protein docking that infers potential interaction by predicting protein complex structure based on the binding energy score[19]; (2) “interaction redundancy” where two structurally similar proteins tend to share the same interaction partners[20]. However, these methods heavily depend on the availability of protein structures, particularly for viral proteins, which remain scarce in databases. In contrast, machine learning-based methods have gained prominence due to their ability to handle large-scale datasets and generate reliable predictions. These approaches leverage machine learning algorithms to extract intrinsic interaction patterns from diverse features, enabling accurate prediction of virus-human PPIs based on the learned patterns. For examples, Yang et al. used doc2vec to encode protein sequences in combination with a Random Forest classifier[21], while Tsukiyama et al. employed word2vec-derived sequence features processed by Long Short-Term Memory (LSTM) networks, followed by a Multilayer Perceptron (MLP) for PPI prediction[22]. Another study by Yang et al. integrated evolutionary sequence profiles with a Siamese Convolutional Neural Network and MLP for PPI prediction[23]. Recently, Zhang et al. used a hybrid attention network to predict virus-human PPIs by integrating protein sequence embeddings from a protein language model and functional annotations[24]. Jiang et al. predicted virus-human PPIs by integrating protein sequence embeddings with a graph neural network and incorporating protein structural information[25]. Bai et al. inferred virus-human PPIs by directly predicting structures of protein complexes[26]. Despite these advances, the field still faces critical challenges, including the lack of standardized benchmark datasets and insufficient consideration of the unique biological characteristics governing virus-human PPIs.

In this study, we firstly constructed a benchmark dataset, ensuring non-overlapped PPIs and minimum sequences similarity of both human and viral proteins in the training and test sets. Then, we proposed vhPPIpred, a novel machine learning-based computational method for virus-human PPI prediction, which extended beyond traditional sequence and phylogenetic features by incorporating human PPIs network topology and viral molecular mimicry of human PPIs. Finally, we applied vhPPIpred to identify human viral receptors. Our results demonstrate that vhPPIpred can serve as an effective tool for predicting virus-human PPIs, thereby advancing the study of viral infection mechanisms and supporting the development of targeted antiviral strategies.

## Results

### Architecture of vhPPIpred

A machine learning-based method named vhPPIpred was developed to predict protein-protein interactions (PPIs) between viruses and human (Fig. 1A). Four kinds of features were incorporated: (1) sequence embedding (a vector of length 1024) generated by ProtT5-XL-U50 that is a pretrained protein language model based on protein sequences of UniRef50[27]; (2) PSSM embedding (a vector of length 20) generated by PSI-BLAST to capture evolutionary information from position-specific scoring matrix (PSSM)[28]; (3) viral molecular mimicry of human protein interactions, which integrated similarities between viral and neighbor proteins that were defined as the proteins interacting with the target protein in the human protein-protein interaction network (HPPIN), and interaction scores between neighbor proteins and the target protein, since viruses can interact with host proteins by mimicking host ligands[29]; (4) degree of human protein in the HPPIN, as proteins with higher degree are more likely targeted by viral proteins[14]. After dimensionality reduction of ProtT5 and PSSM embeddings using principal component analysis (PCA), these four kinds of features were taken as input into several machine learning algorithms for virus-human PPI prediction (see Methods for details).

**Fig. 1.**
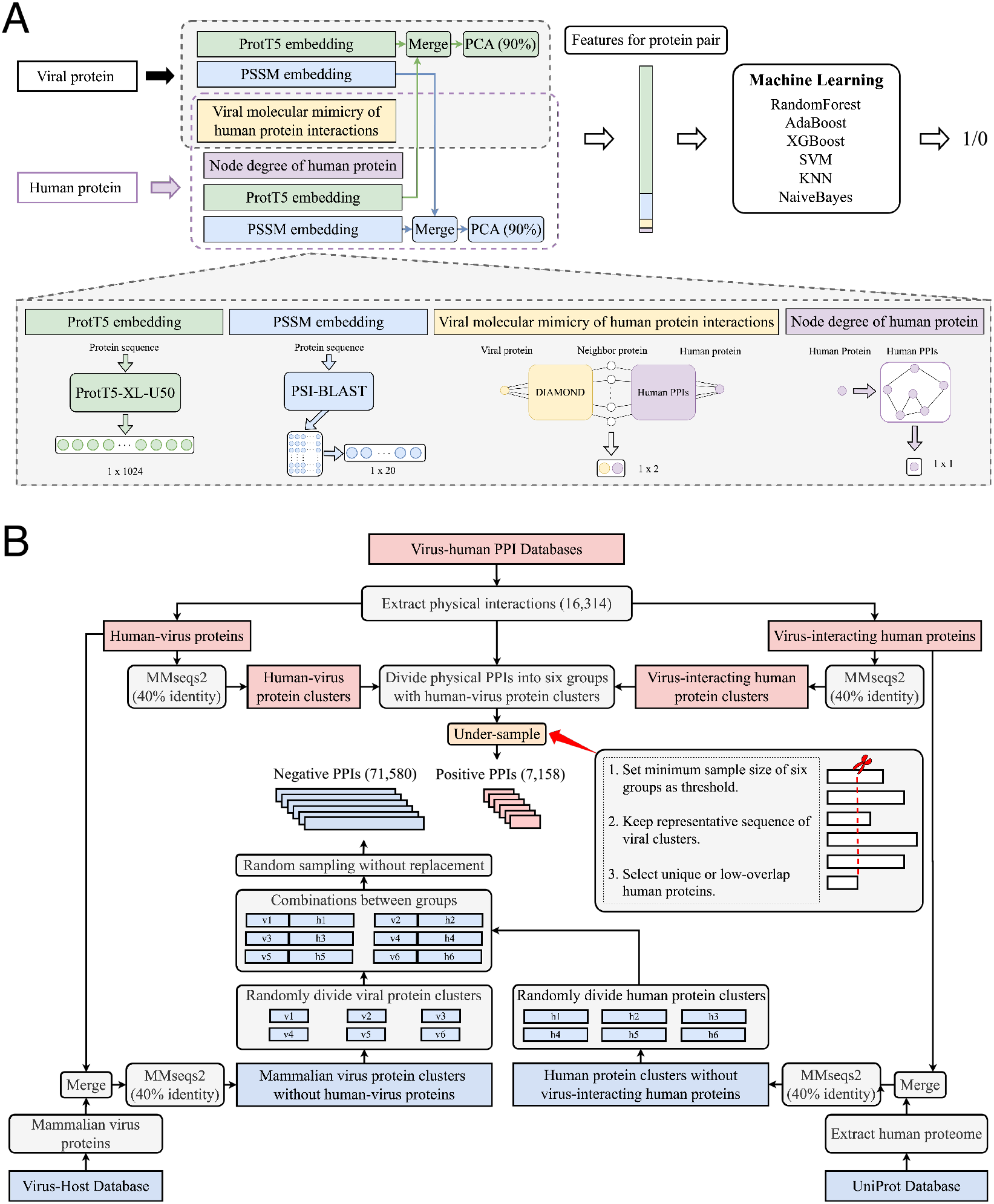
The vhPPIpred architecture and benchmark dataset construction. **(A)** Prediction model integrating multi-source features. **(B)** Stepwise pipeline for benchmark dataset construction.

### Construction of a benchmark dataset

To rigorously evaluate prediction performance, we developed a pipeline to construct a benchmark dataset (Fig. 1B). For positive PPIs construction, we firstly collected 16,314 physical virus-human PPIs involving 938 viral proteins and 5,848 human proteins from eight databases: BioGRID, IntAct, VirusMentha, VirusMINT, VirHostNet, Viruses.STRING, HVIDB, and HVPPI[12–14,30–34]. Then viral and human proteins were independently clustered using MMseqs2[35] at 40% sequence identity, resulting in 501 viral protein clusters and 4,872 human protein clusters. Based on viral protein clusters, all physical PPIs were divided into six groups to ensure viral protein independence across groups. To balance sample sizes, each group was under-sampled to match the smallest group. PPIs involving unique or least-similar human proteins among groups were prioritized to minimize inter-group overlap. In total, 7,158 interactions were retained as positive PPIs through repeated under-sampling.

For negative PPIs construction, we firstly retrieved proteins of viruses known to infect mammals from the Virus-Host Database[36]. These mammalian virus proteins were merged with human-virus proteins and clustered using MMseqs2 at 40% sequence identity. After removing clusters containing any human-virus proteins, we obtained 1,818 viral protein clusters. The human proteome was then downloaded from UniProt[37] and similarly clustered. Clusters containing any virus-interacting human proteins were excluded, resulting in 4,087 human protein clusters. Subsequently, viral and human protein clusters were independently divided into six groups. Negative samples were generated by randomly combining proteins between corresponding viral and human groups, producing millions of candidate interactions. To maintain a 1:10 ratio of positive to negative PPIs, we randomly selected 71,580 negative interactions without replacement from all candidates. After merging positive and negative PPIs, we constructed the final benchmark dataset that comprised six distinct groups.

To assess the overlap of human protein clusters across groups in our benchmark dataset, we calculated Jaccard indices[38] for all group combinations. As shown in Fig. S1, the maximum observed Jaccard index was 0.05, demonstrating a low degree of overlap. This indicated the independence of human protein clusters among groups in our dataset.

### Select XGBoost as the base algorithm

To rigorously evaluate performance of machine learning algorithms, one group (containing 1,193 positive samples and 11,930 negative samples) of benchmark dataset was used as test set, while the remaining five groups were used as training set for 5-fold cross-validation (4 groups for training, 1 group for validation). This procedure was repeated six times, ensuring that each group was used as the test set exactly once.

To select the optimal algorithm, we evaluated six conventional machine learning algorithms and two deep learning algorithms, including Random Forest (RF), Adaptive Boosting (AdaBoost), eXtreme Gradient Boosting (XGBoost), Support Vector Machine (SVM), K-Nearest Neighbors (KNN), Naïve Bayes, Multilayer Perceptron (MLP) and Feature Group Multi-Head Attention model (FeatGroup-MHAttn) using 5-fold cross-validation on training sets[39–41]. For each algorithm, we computed the Area Under the Receiver Operating Characteristic curve (AUROC) and the Area Under the Precision-Recall Curve (AUPRC) independently for each validation fold. The mean AUROC and AUPRC across all folds were reported as the final performance metrics. As shown in Fig. 2A, XGBoost outperformed other algorithms, achieving the highest average AUROC (0.92) and AUPRC (0.67). In comparison, the MLP achieved an AUROC of 0.90 and an AUPRC of 0.63, while FeatGroup-MHAttn achieved an AUROC of 0.89 and an AUPRC of 0.59. These results suggest that, under the current feature representation and training strategy, deep learning-based models did not outperform XGBoost. Therefore, we selected XGBoost as the base algorithm for vhPPIpred and further optimized its hyperparameters via grid search (see Methods). The final parameter settings were provided in Table S1.

**Fig. 2.**
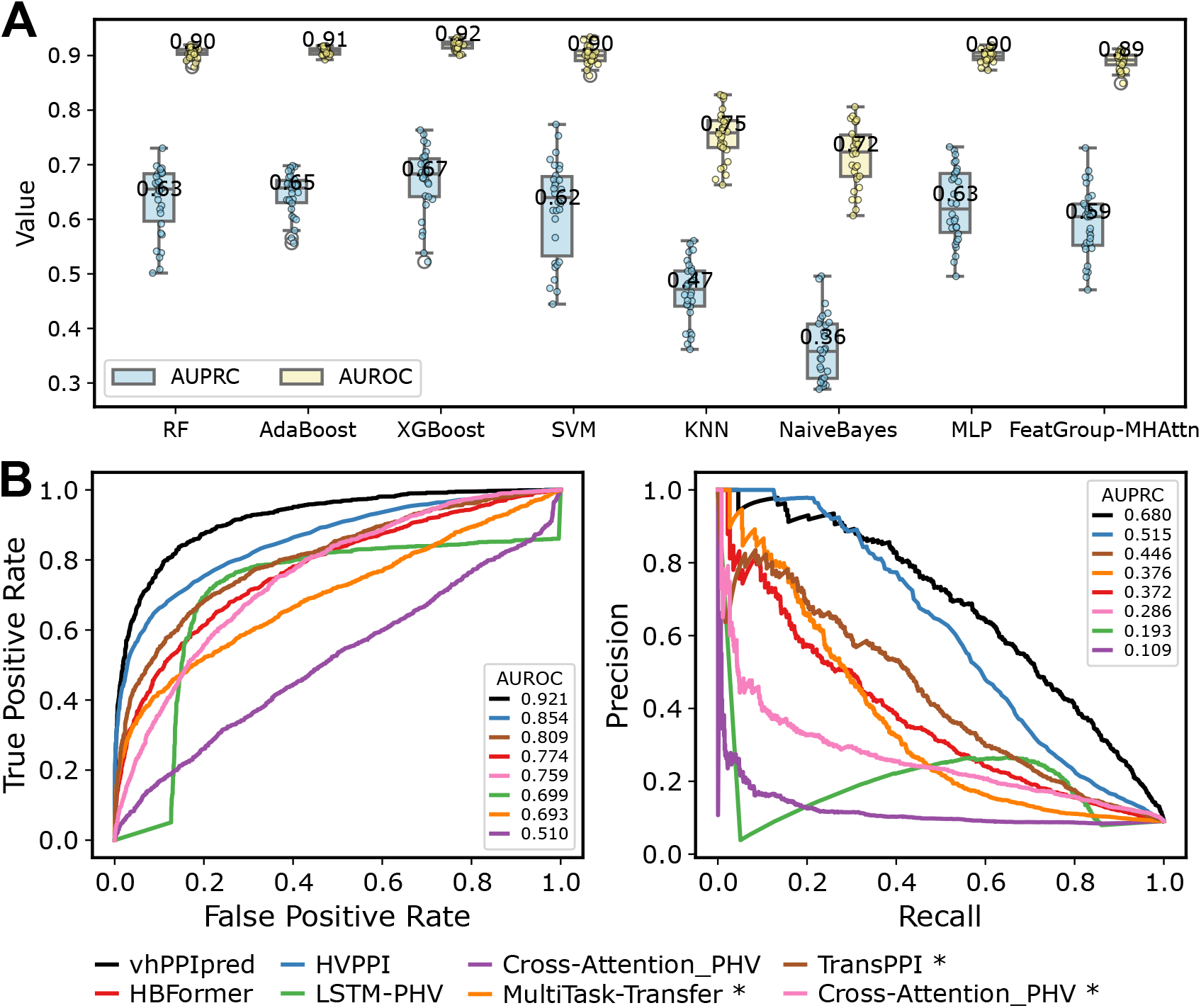
Model construction and evaluation. **(A)** Selection of base prediction algorithm via 5-fold cross-validation on training sets. Boxplots display the distribution of AUROC and AUPRC (averages annotated). **(B)** Performance evaluation and comparison analysis of vhPPIpred against published methods on benchmark dataset. HBFormer, HVPPI, LSTM-PHV and Cross-Attention_PHV were directly tested using their established models. Methods marked with “*” (MultiTask-Transfer, TransPPI and Cross-Attention_PHV) were retrained on our training sets and re-evaluated on corresponding test sets.

### vhPPIpred outperforms other methods on benchmark dataset

To assess the performance of vhPPIpred, we compared it with six published sequence-based and three structure-based virus-human PPI prediction methods on the test set of our benchmark dataset. The sequence-based methods included: (1) HBFormer[24], which employed protein sequence embeddings generated by ProtT5 and predicted virus-human PPIs based on hybrid attention network; (2) HVPPI[21], which employed doc2vec-based protein sequence embeddings and predicted PPIs with a Random Forest classifier; (3) LSTM-PHV[22], which employed word2vec-encoded sequence features processed by Long Short-Term Memory (LSTM) networks and predicted PPIs with Multilayer Perceptron (MLP); (4) Cross-Attention_PHV[42], which captured interaction patterns from word2vec-based protein sequence representations with one-dimensional Convolutional Neural Networks (1D-CNNs) and cross-attention mechanism and predicted PPIs with MLP; (5) MultiTask-Transfer[43], which extracted protein sequence features with Multiplicative LSTM and predicted PPIs with MLP; (6) TransPPI[23], which used 1D-CNNs to extract features from PSSM profiles and predicted interactions with MLP. The structure-based methods included: (1) DeepGNHV[25], which integrated protein sequence embedding and the spatial distribution of amino acids with a graph neural network to generate virus-human PPI representation and predicted interactions using MLP; (2) AlphaFold-multimer[44,45], a derivative version of AlphaFold specifically developed for predicting protein multimer structures; (3) AlphaFold3[46], a diffusion-based model for predicting the joint structures of complexes, including proteins, nucleic acids, and other molecules.

Compared with sequence-based methods, vhPPIpred achieved the highest performance in terms of precision (0.952), AUROC (0.921) and AUPRC (0.680), demonstrating its ability to effectively distinguish between positive and negative samples (Fig. 2B and Table S2). HVPPI showed the second highest AUROC (0.854) and AUPRC (0.515), indicating that it remained a competitive method. HBFormer showed the highest accuracy (0.916) but its recall was relatively low (0.106), suggesting that its high accuracy was mainly driven by correct classification of negative samples rather than efficient detection of positive PPIs. In contrast, LSTM-PHV attained the best recall (0.678) and F1-score (0.378), although its low precision (0.262) resulted in more false positives. Cross-Attention_PHV showed the most constrained performance overall, with particularly poor AUPRC (0.109) that revealed its weakness in recognizing positive interactions. Since the training datasets of these methods partially overlapped with our benchmark testing dataset, their performances might be overestimated. Thus, we removed overlapped PPIs from benchmark dataset and re-evaluated these methods. As shown in Table S2, all methods experienced performance drops across multiple metrics after overlap removal. For HBFormer and HVPPI, although accuracy slightly improved, all other metrics declined. The accuracy of HBFormer improved by 0.033, whereas its AUROC decreased from 0.774 to 0.750 and its AUPRC from 0.372 to 0.251. In contrast, HVPPI showed a larger decrease, with AUROC decreasing by 0.061 (from 0.854 to 0.793) and AUPRC decreasing by 0.299 (from 0.515 to 0.216). LSTM-PHV also declined across all metrics, especially in AUROC (from 0.699 to 0.597) and AUPRC (from 0.193 to 0.107). Cross-Attention_PHV showed a slight accuracy improvement of 0.008, but this was accompanied by declines in other metrics, including AUROC (from 0.510 to 0.492) and AUPRC (from 0.109 to 0.088). These findings collectively suggested that the original performances of these models were likely overestimated due to overlaps between their training and our benchmark testing datasets.

As the trained models of MultiTask-Transfer and TransPPI were not publicly available, we trained and tested them on benchmark dataset using the same training and evaluation protocols as vhPPIpred (see Methods). As shown in Fig. 2B and Table S2, TransPPI outperformed MultiTask-Transfer across all evaluation metrics. However, despite achieving better recall and F1-score, TransPPI performed worse than vhPPIpred in other key metrics, with particularly large gaps in AUROC (a reduction of 0.112) and AUPRC (a reduction of 0.234). Besides, we also retrained and evaluated Cross-Attention_PHV (indicated with “*” in Fig. 2B and Table S2) under the same experimental conditions as vhPPIpred, since difference in the original training datasets might introduce biases into model evaluations. Notably, the retrained model showed improved performance, supporting the notion that its original results were likely influenced by model biases. Nonetheless, the retrained Cross-Attention_PHV still underperformed compared to vhPPIpred, achieving an AUROC of 0.759 and an AUPRC of 0.286, both metrics substantially lower than vhPPIpred’s corresponding values. Additionally, we compared vhPPIpred with three structure-based PPI prediction methods, including DeepGNHV, AlphaFold-multimer, and AlphaFold3. Because the current AlphaFold3 web server only allows a limited number of prediction tasks, we randomly selected 50 positive and 250 negative samples from our benchmark dataset for structure-based performance comparison. AlphaFold-Multimer and AlphaFold3 were directly used to predict virus-human protein complex structures, and the interface confidence score was used as a structure-based proxy score for ranking potential interactions (see Methods). DeepGNHV was directly applied to these samples using its published model. For a fair comparison, vhPPIpred was retrained on the remaining samples after removing the selected samples and those that involve proteins with high sequence similarity to these selected samples (see Methods). As shown in Table S3, vhPPIpred outperformed these structure-based methods across all metrics, especially achieving the highest AUROC (0.972) and AUPRC (0.854). To investigate the reasons underlying the poor performance of structure-based methods, we analyzed the quality of the predicted protein structures involved in the testing samples, including protein monomers and virus-human protein dimers. The results showed that approximately half of the monomers had low confidence scores (Fig. S3A, pLDDT < 70), and most predicted dimers had very low interface confidence scores (Fig. S3B&C, iPTM < 0.6). These low-confidence structures may limit the performance of structure-based methods for large-scale virus-human PPI prediction. In summary, we systematically compared performance of vhPPIpred with nine published methods using a unified benchmark dataset. The results consistently demonstrated that vhPPIpred outperformed other models across most evaluation metrics, highlighting its effectiveness in identifying virus-human PPIs.

### vhPPIpred outperforms other methods on independent datasets

We further compared vhPPIpred with established models of HBFormer, HVPPI, LSTM-PHV and Cross-Attention_PHV on three independent datasets, including Yang’s dataset, Zhou’s dataset and DeNovo dataset[47–49], to evaluate performance and robustness of vhPPIpred.

The first independent test set was Yang’s dataset, comprising 1,241 positive and 12,410 negative PPIs between human and human betaherpesvirus 5 (HHV-5). Because it contains only a single virus type, it primarily evaluates model performance in a virus-specific external setting rather than broad cross-virus generalization. Overlapping PPIs with each method’s training set were removed from Yang’s dataset to prevent data leakage. As shown in Table 1, vhPPIpred achieved the highest F1-score (0.329), AUROC (0.801), and AUPRC (0.275), demonstrating a balanced ability to detect positive interactions while maintaining overall discriminative performance. HBFormer reached the highest precision (0.400) but exhibited very low recall (0.004) and F1-score (0.008), indicating extremely limited sensitivity to positive PPIs. HVPPI achieved the highest accuracy (0.988) but very low precision (0.014) and AUPRC (0.007), suggesting that its high accuracy was largely driven by the abundant negative samples. LSTM-PHV showed the highest recall (0.818) but low precision (0.088) and AUPRC (0.234), reflecting a tendency for false positives. Cross-Attention_PHV displayed lower performance than vhPPIpred across all metrics. Overall, vhPPIpred exhibited the most balanced performance on Yang’s dataset, highlighting its robustness in a virus-specific external evaluation.

**Table 1.**
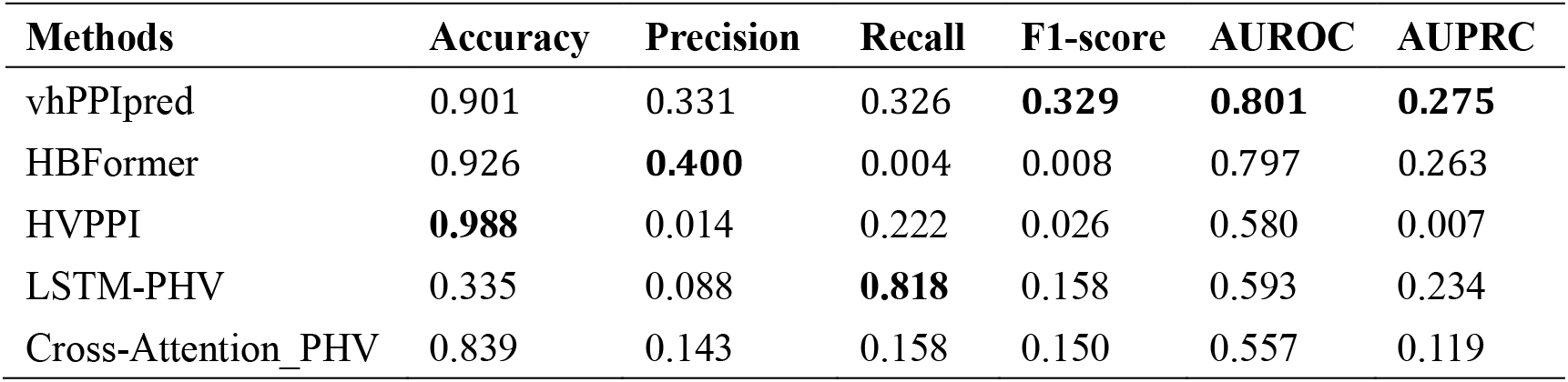
Performance of prediction methods on Yang’s dataset. The evaluated metrics include Accuracy, Precision, Recall, F1-score, AUROC, and AUPRC across different methods. For each metric, the best-performing value is highlighted in bold.

The second independent test set was Zhou’s dataset, containing 749 high-quality binary and co-complex interactions between human and SARS-CoV-2 proteins. Negative samples were generated at ten times the number of positive interactions using the sequence dissimilarity-based strategy[49], and overlapping PPIs with each method’s training set were discarded. Overall, all evaluated methods showed limited predictive performance on this dataset (Table 2). Among them, vhPPIpred achieved the highest recall (0.173) and F1-Score (0.158), indicating relatively better sensitivity to positive PPIs, and also attained the highest AUROC (0.602) and AUPRC (0.179). In contrast, HBFormer reached the highest accuracy (0.917) but exhibited extremely low recall (0.006), and HVPPI showed the highest precision (0.158) with recall of only 0.004, suggesting poor ability to identify positive interactions. LSTM-PHV and Cross-Attention_PHV demonstrated limited performance across all metrics. These results highlight the challenge of predicting PPIs between human proteins and a novel virus, emphasizing the need for further improvements in model design and training.

**Table 2.** Performance of prediction methods on Zhou’s dataset. The best performance of each measure was highlighted in bold.

| Methods | Accuracy | Precision | Recall | F1-Score | AUROC | AUPRC |
| --- | --- | --- | --- | --- | --- | --- |
| vhPPIpred | 0.895 | 0.145 | **0.173** | **0.158** | **0.602** | **0.179** |
| HBFormer | **0.917** | 0.121 | 0.006 | 0.012 | 0.554 | 0.090 |
| HVPPI | 0.908 | **0.158** | 0.004 | 0.008 | 0.521 | 0.100 |
| LSTM-PHV | 0.466 | 0.021 | 0.108 | 0.036 | 0.500 | 0.134 |
| Cross-Attention_PHV | 0.831 | 0.095 | 0.102 | 0.098 | 0.500 | 0.091 |

The third independent test set was the DeNovo dataset, comprising 5,151 PPIs between human and viruses, all curated from the VirusMentha database. Similar to Zhou’s dataset, we constructed negative samples with the same strategy. After removing overlapping PPIs with each method’s training set, all methods were evaluated on the remaining samples. As shown in Table 3, vhPPIpred achieved the highest recall (0.705) and F1-score (0.230), indicating strong sensitivity for detecting true positive interactions, and also attained the highest AUROC (0.698) and AUPRC (0.331), reflecting superior overall discriminative performance. In contrast, HBFormer reached the highest accuracy (0.914) but exhibited extremely low recall (0.008) and F1-score (0.015), suggesting that it largely failed to identify positive PPIs. HVPPI showed high accuracy (0.961) but a low F1-score (0.124), moderate recall (0.229) and limited precision (0.085). LSTM-PHV and Cross-Attention_PHV demonstrated intermediate performance across all metrics, with lower AUROC and AUPRC values (0.546, 0.549 and 0.115, 0.177, respectively), reflecting their restricted predictive ability on this dataset. Overall, these results highlight that vhPPIpred offers the best balance between positive-sample detection and overall classification performance on the DeNovo dataset.

**Table 3.** Performance of prediction methods on DeNovo dataset. The best performance of each measure was highlighted in bold.

| Methods | Accuracy | Precision | Recall | F1-Score | AUROC | AUPRC |
| --- | --- | --- | --- | --- | --- | --- |
| vhPPIpred | 0.576 | 0.137 | **0.705** | **0.230** | **0.698** | **0.331** |
| HBFormer | 0.914 | **0.275** | 0.008 | 0.015 | 0.627 | 0.133 |
| HVPPI | **0.961** | 0.085 | 0.229 | 0.124 | 0.692 | 0.075 |
| LSTM-PHV | 0.869 | 0.209 | 0.157 | 0.179 | 0.546 | 0.177 |
| Cross-Attention_PHV | 0.833 | 0.147 | 0.174 | 0.159 | 0.549 | 0.115 |

In summary, vhPPIpred consistently outperformed the other methods on these three independent test sets. These results demonstrate its relative robustness and generalization ability across different virus-human interaction scenarios, underscoring its potential for practical applications.

### Feature contribution analysis of vhPPIpred

To evaluate the contribution of each feature type in vhPPIpred, we conducted a systematic analysis by testing all possible combinations of the four feature categories. As shown in Fig. 3A, when the feature type was used alone, the model using ProtT5 embeddings achieved the highest AUROC of 0.872, whereas the viral molecular mimicry feature achieved the highest AUPRC of 0.562. Among the pairwise feature combinations, the combination of ProtT5 embeddings and human protein degree markedly improved model performance, with an AUROC over 0.90 and an AUPRC over 0.65. Based on ProtT5 embeddings and human protein degree, adding either viral molecular mimicry or PSSM embeddings resulted in similar performance, with the highest AUROC of 0.917 and AUPRC value of 0.677, respectively. The model integrating all four feature types achieved the best overall performance, with the highest AUROC of 0.921 and AUPRC of 0.680. These results indicate that ProtT5 embeddings and human protein degree contributed more strongly to vhPPIpred than viral molecular mimicry and PSSM embeddings, while the latter two features still provided modest complementary information.

**Fig. 3.**
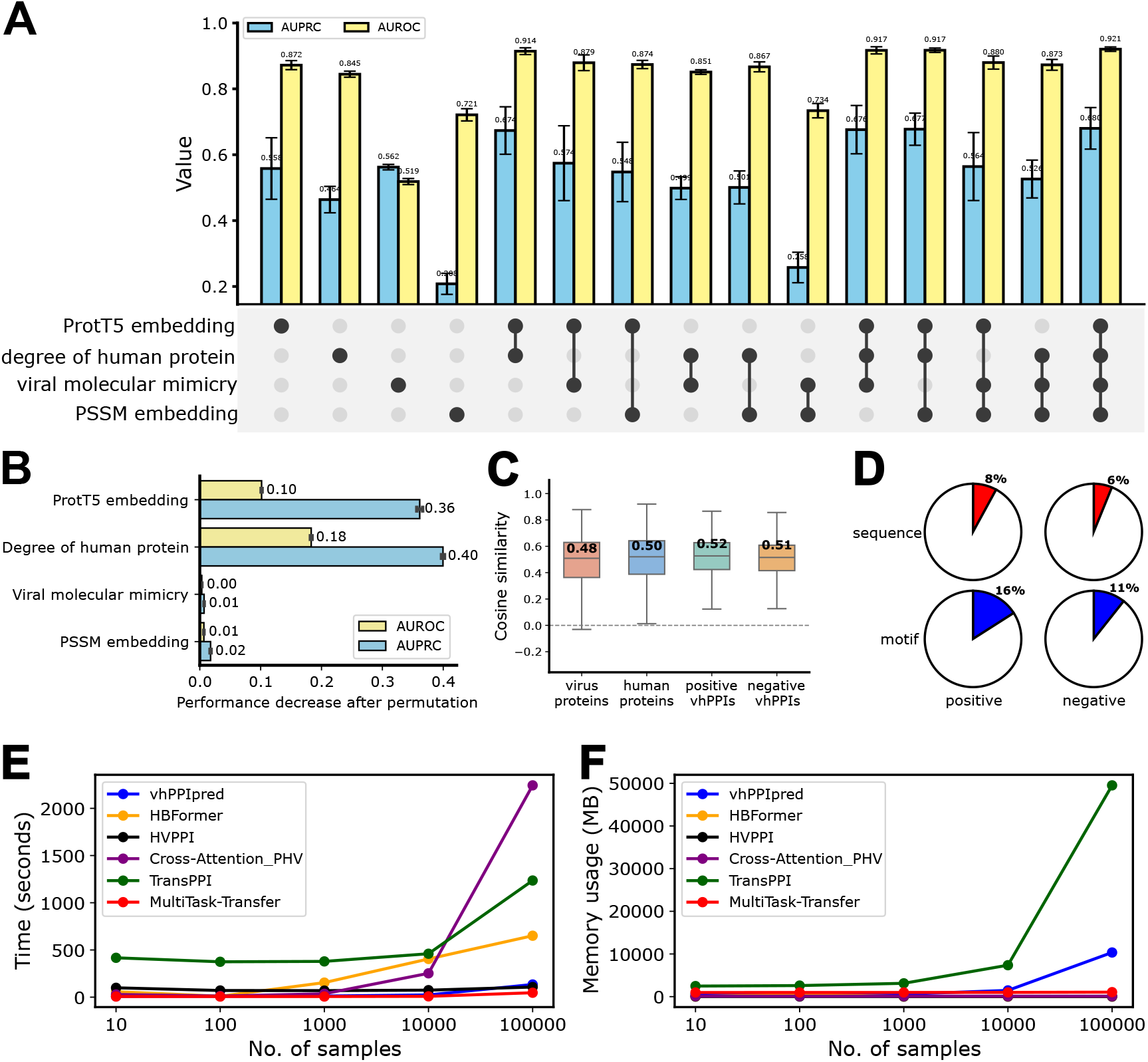
Feature contribution analysis of vhPPIpred and time-space complexity of multiple prediction methods. **(A)** Performance of vhPPIpred using all combinations of the four feature types: ProtT5 embeddings, degree of human protein, viral molecular mimicry and PSSM embeddings. **(B)** Performance decrease after permutating each feature type. **(C)** Cosine similarity between ProtT5 embeddings and PSSM embeddings for virus proteins, human protein, positive virus-human PPIs, and negative virus-human PPIs. **(D)** Coverage of positive and negative samples with similar sequences or motifs between viral proteins and neighbor proteins. **(E)** Time consumption measured on datasets of varying sample sizes. **(F)** Memory usage measured during predictions on datasets of varying sample sizes. All methods were performed on Ubuntu 18.04.2 LTS system. The CPU used was an Intel(R) Xeon(R) E5-2698 v4 CPU @ 2.20GHz, with 84 GB of memory.

We further performed additional analyses to improve model interpretability. As shown in Fig. 3B, permutation-based feature importance analysis demonstrated that permuting ProtT5 embeddings or human protein degree caused relatively large decreases in model performance, with AUROC decreases of 0.10 and 0.18 and AUPRC decreases of 0.36 and 0.40, respectively. In contrast, permuting viral molecular mimicry or PSSM embeddings resulted in only minimal performance changes, suggesting that these two features contributed relatively limited additional information under the current feature design. These findings suggest that vhPPIpred mainly relies on protein language model-derived representations and host protein network-related information, whereas molecular mimicry and PSSM provide relatively limited additional information under the current feature design.

To explain the limited contribution of PSSM embeddings, we calculated the cosine similarity between ProtT5 and PSSM embeddings at both protein and virus-human PPI (vhPPI) levels (see Methods for details). As shown in Fig. 3C, the median similarities were 0.48 for viral proteins, 0.50 for human proteins, 0.52 for positive vhPPIs, and 0.51 for negative vhPPIs, indicating a moderate overlap between ProtT5 and PSSM representations. This overlap suggests that PSSM embeddings share part of the sequence- and evolution-related information with the ProtT5 embeddings. Although PSSM embeddings may still contain some residual information not fully represented by ProtT5, this additional information appears to provide only limited incremental value for vhPPI prediction. Therefore, the redundancy between PSSM and ProtT5 embeddings may explain the small performance gain observed after adding PSSM features.

To further investigate the limited contribution of viral molecular mimicry feature, we calculated the sequence similarity and motif-level similarity between the viral proteins and the neighbor proteins (defined as proteins that interact with virus-infecting human proteins) (see Methods). As shown in Fig. 3D, only a small fraction of protein pairs exhibited sequence similarity, with positive pairs slightly higher than negative ones (8% vs 6%). Similar results were observed at the motif level, although the proportion of motif-level similarity (16% and 11%) was higher than that at the sequence level. These observations indicate that although viral molecular mimicry at both the sequence and motif level is more frequently detected in positive samples than in negative pairs, supporting its biological relevance, the sequence or motif similarity is only present in a subset of pairs, which may explain the minor contribution of viral molecular mimicry feature in predicting virus-human PPIs.

### Analysis of time consumption and memory usage

To quantify computational resource usage, we randomly took N (N = 10, 100, 1,000, 10,000, 100,000) samples from our benchmark dataset and measured the execution time and memory usage of each method across different sample sizes in predicting interactions (Fig. 3E&F). For time consumption, it was evident that most methods exhibited relatively stable runtimes at small to moderate sample sizes. However, significant differences emerged as the number of samples increased. HBFormer, TransPPI and Cross-Attention_PHV showed sharp increases in execution time when processing 10,000 or more samples, with Cross-Attention_PHV being the most affected. In contrast, vhPPIpred, HVPPI and MultiTask-Transfer displayed a more gradual increase, indicating their computational efficiency in handling large-scale data.

For memory usage, most methods maintained relatively low and stable memory consumption at small to moderate sample sizes. HBFormer, HVPPI, Cross-Attention_PHV, and MultiTask-Transfer maintained stable memory requirements, indicating their suitability for large-scale datasets in terms of memory efficiency. However, as the number of samples approached 100,000, TransPPI exhibited a steep increase in memory consumption, surpassing all other methods. vhPPIpred also showed a notable rise, though to a lesser extent.

### Application of vhPPIpred in identification of viral receptors

Host receptors are essential for viral entry, yet their identification remains challenging. To evaluate the ability of computational methods in predicting viral receptors, we applied vhPPIpred, HBFormer, HVPPI, LSTM-PHV and Cross-Attention_PHV to predict interactions between viral receptor binding proteins (RBPs) and human receptors, since these methods provided publicly available models. We firstly downloaded 213 known RBP-receptor pairs (involving 103 RBPs and 87 human receptors) from the Viral Receptor database[50]. Then we collected 1,418 potential receptor candidates of human-infecting virome from Zhang’s dataset[51]. For each RBP, we predicted interaction probabilities with all potential receptors. We subsequently sorted potential receptors with predicted confidence scores in descending order and recorded ranks of true receptors. After predicting interactions between all RBPs and all potential receptors, we obtained ranks of true receptors in all RBPs. Finally, we quantified each method’s performance by counting how many true receptors were ranked within the top N (N=10, 50) predictions.

As shown in Fig. 4A, vhPPIpred identified 7 known RBP-receptor pairs within the top 10 predictions, whereas HBFormer and Cross-Attention_PHV each identified one pair, and HVPPI and LSTM-PHV identified none. Within the top 50 predictions, both vhPPIpred and HBFomer identified18 known pairs identified, followed by LSTM-PHV, which identified 13 known pairs. These results suggest that vhPPIpred showed favorable performance in prioritizing known viral receptors, especially among the top-ranked predictions.

**Fig. 4.**
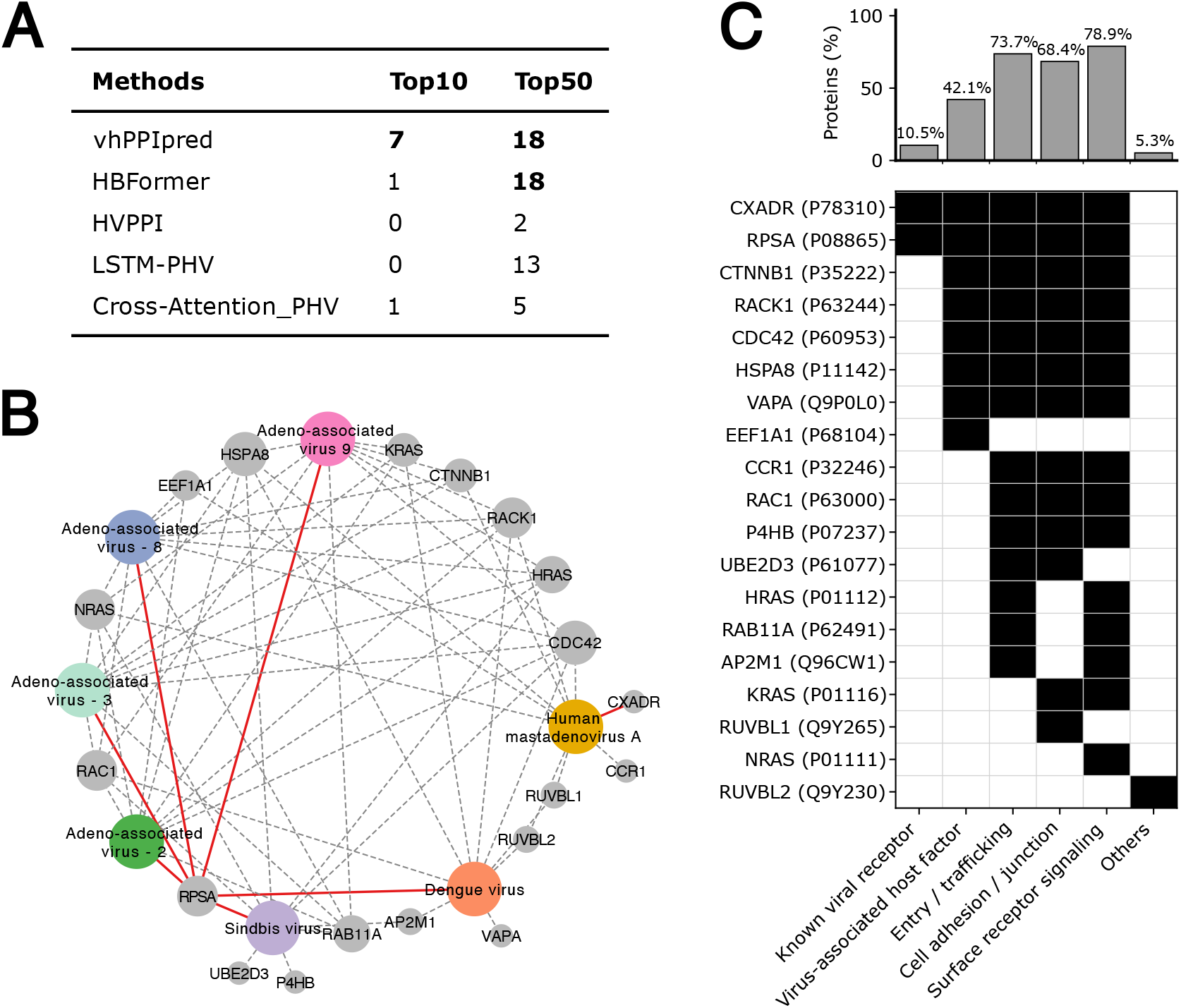
Performance of prediction methods on identifying viral receptors and qualitative analysis of receptor candidates identified by vhPPIpred. **(A)** The number of known receptors identified by different methods in the Top10 and Top50 predictions is reported, with the highest value for each method highlighted in bold. **(B)** Network visualization of the top 10 predicted human genes for seven viruses whose known receptors were successfully identified by vhPPIpred. Viruses are shown as colored nodes, whereas human genes are shown as gray nodes. Node size is proportional to the degree in the prediction network. Red solid edges indicate experimentally validated virus–receptor interactions (known receptor pairs), while gray dashed edges indicate high-ranking label-negative predicted candidates. **(C)** Qualitative functional classification of the top-ranked predicted human candidates based on curated UniProt annotations. The upper bar plot shows the proportion of human genes assigned to each functional category. The lower binary matrix indicates the presence (black) or absence (white) of corresponding functional evidence for each protein.

To further interpret viral receptor predictions of vhPPIpred, we performed a qualitative analysis of the top 10 predictions for seven viral proteins whose known receptors were successfully ranked within the top 10 predictions. This analysis resulted in 70 predicted virus-human protein pairs involving 19 human cell membrane genes (Fig. 4B). Among these human genes, two known viral receptors, RPSA and CXADR, were successfully identified by vhPPIpred. Annotations of these top-ranked human genes were manually curated from the UniProt database and classified into six categories: known viral receptor, virus-associated host factor, entry/trafficking-related protein, cell adhesion/junction-associated protein, surface receptor signaling protein, and others. As shown in Fig. 4C, most of these top-ranked genes were associated with biological processes relevant to viral infections. For example, 12 genes were annotated as surface receptor signaling or cell adhesion/junction, which may participate in viral attachment, entry, or host cell interaction processes. Therefore, these label-negative high-ranking predictions should not be simply interpreted as definitive false positives. Instead, some of them may represent biologically plausible but currently unannotated candidate receptors.

## Discussion

In this work, we constructed a benchmark dataset for rigorous evaluation of methods in predicting virus-human PPIs. Then we developed a machine learning-based prediction method, named vhPPIpred, which integrated multi-source features to improve the prediction of virus-human PPIs. Comparative experiments showed that vhPPIpred consistently outperformed existing methods on our benchmark dataset and three independent datasets. Subsequently, it was applied to the identification of human viral receptors, showing promising potential for receptor prediction. Feature contribution analysis highlighted the value of integrating multiple features for PPI prediction. Time and space complexity analysis further revealed that vhPPIpred required fewer computational resources, suggesting its broad applicability in virus-human PPI prediction.

There are two fundamental limitations in existing virus-human PPI prediction methods. The first is the lack of benchmark datasets, particularly due to the absence of biologically confirmed negative samples. Current negative sampling strategies typically fall into two categories: (1) random sampling, where negative samples are generated by random pairing viral and human proteins from positive samples while excluding known interactions[52]; (2) dissimilarity-based sampling, which further eliminates protein pairs with sequence similarity to positives[49]. However, both approaches may inadvertently introduce false negatives, potentially compromising model reliability. To mitigate this risk, we exclusively employ mammalian virus proteins and remove sequences showing similarity to known human virus proteins. This dual consideration of biological origin and sequence dissimilarity provides a more reliable foundation for constructing negative samples with reduced risk of containing true interactions. While this strategy greatly minimizes such risk, we acknowledge that the inherent complexity of protein interactions makes it impossible to eliminate all false negatives.

The second limitation is the potential overestimation of performance due to inappropriate dataset partitioning. Existing prediction methods typically divide dataset randomly into training and test set[21,22,42]. However, this strategy often introduces overlapped proteins between training and test set, which can result in overestimation of prediction performance due to information leakage. To address this issue, we separately divided positive and negative samples into six distinct groups based on viral protein clusters. For each cluster, we selected representative viral proteins and ensured that human proteins were mutually dissimilar across groups (as shown in Fig. S1). This design ensured non-overlapped PPIs and minimum sequences similarity of both human and viral proteins in the training and test sets, thus enabling a more rigorous and unbiased evaluation of predictive performance. More importantly, the benchmark dataset can serve as a standardized and biologically meaningful resource for future research on virus-human PPI prediction.

Accurate protein representation is critical for predicting virus-human PPIs. Current methods typically rely on amino acid composition, physicochemical properties, sequence embeddings, or evolutionary information from protein sequences[53–55]. However, these approaches often overlook the intrinsic characteristics of interactions and may fail to capture the complexity of virus-host recognition mechanisms. To address these limitations, we developed vhPPIpred that predicted virus-human PPIs by integrating four kinds of features: (1) protein sequence features, (2) evolution features, (3) human protein degree, and (4) viral molecular mimicry of human protein interactions. Sequence and evolution features have been widely used in existing methods, underscoring their foundational role in protein representation. More importantly, our method incorporates two key biological characteristics of virus-host interactions: (1) viral proteins interact with host proteins by mimicking host ligands[29], and (2) viral proteins preferentially target human proteins with higher node degrees in the human PPI network[14]. These features focus on interaction-specific attributes, rather than relying solely on intrinsic protein properties.

Feature contribution analysis showed that ProtT5 embeddings and human protein degree contributed more strongly to model performance than viral molecular mimicry and PSSM embeddings. Nevertheless, the integration of all four feature types achieved the best overall performance, suggesting that viral molecular mimicry and PSSM embeddings still provide complementary information. Additional interpretability analyses further explained why these two features showed relatively limited contributions. The weak contribution of PSSM embeddings may be due to redundancy with ProtT5 embeddings, as protein language models can encode evolutionary information that overlaps with PSSM-derived features. Additionally, the viral molecular mimicry feature based on sequence similarity is likely limited by the restricted coverage of sequence similarity in capturing short motif-level mimicry that is important for virus-human PPIs[56–58]. These results indicate that incorporating more detailed motif-level representation, such as context-specific or structural motif representations, to increase the coverage of sequence similarity searches may further improve virus-human PPI prediction.

Our method also has potential utility in biological studies. We showed that vhPPIpred could be used to prioritize candidate human membrane proteins for viral receptor prediction. Compared with other virus-human PPI prediction methods, vhPPIpred showed favorable recovery of known RBP-receptor pairs among top-ranked predictions. Importantly, qualitative analysis suggested that several label-negative high-ranking predictions had functional annotations related to viral entry or infection-associated host processes. This observation highlights the incompleteness of current viral receptor annotations and suggests that some top-ranked label-negative predictions may represent biologically plausible candidate receptors or virus-associated host factors. Nevertheless, these predictions should be regarded as computational candidates and require further experimental validation before being considered as real viral receptors.

While vhPPIpred represents significant advances in predicting virus-human PPIs, several limitations should be considered. Firstly, although we constructed negative samples using mammalian virus proteins, undiscovered interactions between these viral proteins and human proteins could lead to false negatives. This limitation might be addressed by adopting strategies similar to the Negatome Database[59], which systematically derived non-interacting protein pairs through literature mining, manual annotation and structural analysis of protein interfaces. Secondly, although ProtT5-XL-U50 achieved the best performance among the evaluated protein language models (ProtBert and ESM-2, Table S4), the rapid development of pretrained protein language models suggests that larger or more recent models, such as ProtT5-XXL-U50 and ESM-3, may provide improved protein representations in future studies[27,60,61]. Thirdly, although vhPPIpred showed favorable performance across multiple benchmark datasets, its sensitivity was reduced in the challenging SARS-CoV-2-specific Zhou’s dataset, similar to other competing methods. This suggests that the generalization ability of vhPPI predictors for highly divergent viruses, newly emerging viruses, or virus-specific benchmark scenarios still requires further improvement. Finally, similar to several existing approaches, vhPPIpred currently does not incorporate structural information due to the limited availability of high-quality viral protein structures. Although databases such as the Big Fantastic Virus Database (BFVD)[62] and Viro3D[63] provided some viral protein structures, the quality of most remains insufficient for reliable modelling. However, some methods might improve predictions of viral protein structures. For instance, DeepMSA2[64] can construct multiple-sequence alignment profile containing richer evolutionary information through iterative alignment search across diverse sequence databases, while ProteinTTT[65] can provide protein-specific representation by fine-tuning a pretrained protein language model on one protein sequence.

### Conclusions

This study provides a valuable benchmark dataset for developing and evaluation of virus-human PPI prediction methods. A state-of-the-art method named vhPPIpred was further developed to predict virus-human PPIs and outperformed other methods on both the benchmark dataset and three independent datasets. The method was demonstrated to be useful in identification of viral receptors. The study advances computational approaches for virus-human PPIs identification, with potential applications in antiviral drug discovery and virus-human interaction research.

## Methods

### Integration of virus-human protein-protein interactions

We downloaded protein-protein interactions (PPIs) between viruses and human from eight databases: BioGRID (v4.4.208)[30], IntAct (v4.2.3.2)[31], VirusMentha (v1.0)[32], VirusMINT (v1.0)[33], VirHostNet (v3.0)[34], Viruses.STRING (v10.5)[12], HVIDB (v1.0)[13] and HVPPI (v1.0)[14] on May 16, 2022. To ensure consistency across datasets, we first standardized the protein identifiers by converting various ID types (including gene symbols, Ensembl IDs) into UniProt accession numbers using bioDBnet (v2.1, https://biodbnet-abcc.ncifcrf.gov/db/db2db.php)[66]. Then, we applied max-min normalization to interaction scores within each dataset to make scores comparable across databases. After integrating all normalized virus-human PPIs, we removed duplicates by retaining only the highest-scoring entry for each unique interaction pair within the same interaction type, resulting in a comprehensive dataset of 108,331 interaction pairs.

To enhance data reliability, we filtered interactions by selecting those annotated with direct physical interaction keywords (“physical association”, “direct interaction”, “physical” or “covalent binding”). For interactions associated with multiple keywords, only the highest-scoring entry was kept. To exclude viral polyproteins, we retained interactions involving proteins with lengths between 50 and 1,000 amino acids. These steps resulted in a high-confidence dataset of 16,314 interaction pairs, involving 938 viral proteins and 5,848 human proteins.

### Construction of human protein-protein interaction network

As shown in Table S5, we collected human PPIs from six databases. Mentha (v1.0, April 25, 2022)[67] and BioPlex (v3.0, May 30, 2022)[68] provided complete sets of physical PPIs, which were used without additional filtering. For STRING (v11.5, May 16, 2022)[69], we extracted high-confidence physical interactions (combined_score ≥ 900) from the file *9606.protein.physical.links.v11.5.txt.gz* and converted Ensembl IDs to UniProt accession numbers using bioDBnet. For MINT (v1.0, May 5, 2022)[70], IntAct (v4.2.3.2, May 5, 2022)[31] and BioGRID (v4.4.208, June 30, 2022)[30], we retained only interactions annotated with specific binding types.

To ensure score comparability across databases, we applied max-min normalization separately to the interaction scores within each dataset. All normalized human PPIs were then merged, and for interactions reported by multiple databases, only the entry with the highest normalized score was retained. This stringent normalization and deduplication process resulted in a final dataset comprising 605,507 non-redundant human PPIs.

### Collections of mammalian virus proteins

We collected 255 virus species known to infect seven mammalian hosts (*Bos taurus, Canis lupus familiaris, Chlorocebus aethiops, Macaca mulatta, Mesocricetus auratus, Mus musculus, and Sus scrofa*) from the Virus-Host Database (release 217, accessed May 25, 2023)[36]. Importantly, none of these viruses are known to infect human. From this database, we retrieved 5,914 proteins encoded by these viruses. To exclude polyproteins, we filtered the sequences by lengths, keeping only those between 50 and 1,000 amino acids. This resulted in a final dataset of 5,302 mammalian virus proteins.

### Collections of human proteins

The reviewed human proteome was downloaded from the UniProt database (release-2023_01)[37] on April 23, 2023, resulting in an initial dataset of 20,422 proteins. After removing proteins shorter than 50 amino acids, 20,325 human proteins were retained.

### Clustering proteins with MMseq2

MMseqs2 (v13-45111)[35] was downloaded from https://github.com/soedinglab/MMseqs2. We used it to perform sequence clustering of viral and human proteins with a sequence identity threshold of 40% (min-seq-id=0.4).

### Construction of benchmark dataset

The positive samples were constructed based on physical interactions between viral and human proteins. The viral and human proteins from 16,314 high-confidence physical PPIs were clustered separately using MMseqs2 (v13-45111) with a sequence identity threshold of 40%, resulting in 501 viral protein clusters and 4,872 human protein clusters, respectively.

Next, we randomly divided all high-confidence physical interactions into six groups according to viral protein clusters, with each group containing approximately 83 clusters. To ensure consistency in sample size across groups, we determined the size of the smallest group as the threshold and performed under-sampling on the remaining groups to match this size.

In the under-sampling process, for each viral cluster, a representative protein was selected. To minimize the overlap of human proteins between groups, we prioritized the use of unique human proteins that only appeared in a single group. If the number of interactions after under-sampling with unique human proteins was insufficient, we additionally included shared human proteins with low inter-group redundancy. Specifically, we computed the Jaccard indices between the human protein clusters of each group and the union of human clusters from the remaining five groups. Shared human proteins were ranked in ascending order of Jaccard indices and incrementally selected to construct positive interactions. Virus-human PPIs involving representative viral proteins and the selected human proteins were randomly under-sampled without replacement until the number of interactions in each group met the threshold. This procedure resulted in a total of 7,158 positive interactions, with each group containing exactly 1,193 interactions.

To construct negative samples, we firstly merged human virus proteins with mammalian virus proteins and clustered them using MMseqs2 with a sequence identity threshold of 40%. Clusters containing any human virus proteins were removed, resulting in 1,818 mammalian virus protein clusters. Separately, the human proteome was clustered using MMseqs2, and clusters containing any virus-infecting human proteins were excluded, resulting in 4,087 human protein clusters. The remaining mammalian virus and human protein clusters were then randomly divided into six groups, respectively. Negative interaction pairs were generated by randomly combining clusters of viral groups and human groups (e.g., v1-h1, v2-h2, etc., as shown in Fig. 1B). For each group, we sampled negative pairs without replacement, ensuring that the number of negative samples was exactly 10 times the number of corresponding positive samples. This process resulted in 71,580 non-interacting protein pairs, with each group containing exactly 11,930 negative samples.

After integrating the positive and negative samples, we constructed a final benchmark dataset comprising six groups, with a total of 78,738 samples. We adopted an iterative evaluation strategy in which one group was used as the test set and the remaining five groups were used for 5-fold cross-validation (4 groups for training and 1 group for validation in each fold). This process was repeated six times, ensuring that each group was used as the test set exactly once.

To evaluate the overlap of human proteins across groups, we calculated the pairwise Jaccard indices[38] between each of the six groups based on their corresponding human protein clusters (as shown in Fig. S1). The Jaccard index is defined as

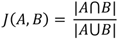

where *A* and *B* are finite sets, |*A*⋂*B*| denotes the cardinality of their intersection, and |*A*⋃*B*| donates the cardinality of their union.

### Yang’s dataset

The Yang’s dataset[47], constructed by Yang et al., contains 1,241 positive and 12,410 negative PPIs between human and human betaherpesvirus 5 (HHV-5). We obtained this dataset from the supplementary materials of the corresponding publication on August 4, 2024.

### Zhou’s dataset

The Zhou’s dataset[48], constructed by Zhou et al., contains 739 gene-gene interactions between human and Severe Acute Respiratory Syndrome coronavirus 2 (SARS-CoV-2). We obtained this dataset from the supplementary materials of the corresponding publication on August 4, 2024. After manually converting gene symbols to protein identifiers (UniProt or NCBI Protein accession numbers), we derived a total of 749 positive protein-protein interactions.

To construct negative virus-human PPI samples, we used a sequence dissimilarity-based negative sampling strategy following the method described by Eid et al[49]. As shown in Fig. S4, we firstly generated all possible pairwise combinations between viral proteins and human proteins observed in positive virus-human PPIs. Viral and human proteins were then separately clustered using MMseqs2 with a sequence identity threshold of 0.4, resulting in viral protein clusters and human protein clusters. Known positive PPIs were directly removed from all possible pairwise combinations. Next, both the remaining protein pairs and the positive PPIs were mapped to the representative proteins of their corresponding viral and human protein clusters, generating representative protein-pair combinations and representative positive PPIs. Representative combinations that overlapped with representative positive PPIs were further removed to exclude protein pairs with high sequence similarity to known positives. The remaining representative combinations were then mapped back to their corresponding original protein pairs to generate a pool of candidate negative PPIs. Finally, negative samples were randomly selected from this pool, with the number of negative PPIs set to ten times that of positive PPIs.

### DeNovo dataset

The DeNovo dataset[49], constructed by Eid et al., contains 5,445 PPIs between human and viruses collected from the VirusMentha database (accessed in June 2014). We downloaded this dataset from the supplementary materials of the corresponding publication on August 4, 2024. After excluding proteins not present in the UniProt database (release-2024_03), we retained a total of 5,151 positive virus-human PPIs. The negative PPIs were constructed using the same strategy as that used for the negative samples in Zhou’s dataset.

### ProtT5 embedding

We downloaded the pretrained language model ProtT5-XL-U50[27] (based on UniRef50) from Hugging Face (https://huggingface.co/Rostlab/prot_t5_xl_half_uniref50-enc) on November 19, 2023. Additionally, we also downloaded another two models, ProtBert[27] and ESM-2[60], from https://huggingface.co/Rostlab/prot_bert and https://github.com/facebookresearch/esm, respectively, on May 19, 2026. After comparing performance integrating protein embeddings generated by different protein language models (Table S4), we finally selected ProtT5-XL-U50 to encode all viral and human protein sequences, generating a vector of length 1,024 for each protein.

### PSSM embedding

To capture evolutionary information from protein sequences, we generated position-specific scoring matrix (PSSM) using PSI-BLAST (NCBI BLAST 2.12.0+)[28]. All viral and human protein sequences were queried against the UniRef90 database[37] (retrieved from UniProt on October 18, 2023) using three iterations (num_iteractions=3). For each protein, a PSSM was generated and then subjected to row-wise normalization. Subsequently, we performed column-wise averaging on each normalized PSSM to obtain a 20-dimensional feature vector per protein.

### PCA of ProtT5 embeddings and PSSM embeddings

To reduce computational resource consumption, we applied principal component analysis (PCA) separately to the ProtT5 and PSSM embeddings at PPI level. Specifically, for each virus-human protein pair, we concatenated the ProtT5 embeddings of viral and human proteins (each 1,024-dimentional) to form a 2,048-dimentional vector. This resulted in an *N* × 2048 matrix, where *N* denotes the number of PPIs. PCA was then performed on this matrix using the decomposition.PCA function from scikit-learn package (v1.3.2)[39], retaining 90% of the variance (n_components=0.9). The PCA was fitted on the training set via the fit_transform method and applied to the test set using the transform method. The same procedure was applied to the PSSM embeddings: we concatenated the 20-dimensional PSSM vectors of each virus–human protein pair to obtain an *N* × 40 matrix and reduced its dimensionality while preserving 90% of the variance.

### Degree of human protein

We utilized NetworkX (v3.1)[71] in Python (v3.8.19)[72] to calculate the degree of each human protein node within the human protein-protein interaction network. The network was represented as an undirected graph *G =* (*V, E*), where the nodes (*V*) represented human proteins and the edges (*E*) represented interactions between them. The degree of each protein was computed using *G*.degree function, which returned the number of direct interaction partners it had.

### Quantify viral molecular mimicry of human protein interactions

Viruses can interact with host proteins by mimicking host ligands[29]. For instance, in the case of viral protein X and human protein A, if X is similar with human proteins that directly interact with A, then X may interact with A. All human proteins directly interacting with A in the human PPI network were defined as neighbor proteins of A. To quantify this viral molecular mimicry, we used DIAMOND blastp (v2.1.9.163)[73] with an e-value threshold of 1E-6 (all other parameters set to default) to compute sequence similarity (bit score) between viral protein X and neighbor proteins of A, averaging the similarity scores when multiple neighbors existed. In parallel, we integrated the normalized interaction scores between neighbor proteins and human protein A, also averaging these scores when multiple neighbors existed. These two values were combined to provide a quantitative measure of viral molecular mimicry of human protein interactions.

### Machine learning algorithms used in prediction of virus-human PPIs

During base algorithm selection, we systematically evaluated six machine learning algorithms (Random Forest, AdaBoost, XGBoost, SVM, Naïve Bayes and KNN)[39,40] and two deep learning models (Multilayer Perceptron and Feature Group Multi-Head Attention model) by tuning key hyperparameters and comparing their performance. The first was the Random Forest classifier, which was implemented using the ensemble.RandomForestClassifier function from the scikit-learn package (v1.3.2) in Python (v3.8.19). For this algorithm, we specifically tuned the number of trees (n_estimators) with values of 50, 100, 300, and 500, while keeping all other parameters at their default settings. Based on performance evaluations, we selected n_estimators=500 as the optimal configuration.

The second was the AdaBoost classifier, which was implemented using the ensemble.AdaBoostClassifier function from the scikit-learn package (v1.3.2) in Python (v3.8.19). For this algorithm, we performed a grid search over key hyperparameters using itertools.product function, evaluating combinations of n_estimators (50, 100, 300, 500) and learning_rate (0.01, 0.1, 1.0), while keeping other parameters at their default settings. Based on performance comparison, we selected n_estimators=300 and learning_rate=0.1 as the optimal configuration.

The third was the XGBoost classifier, which was implemented using the XGBClassifier function from the XGBoost package (v2.1.1) in Python (v3.8.19). For this algorithm, we conducted an exhaustive grid search over n_estimators (50, 100, 300, 500) and learning_rate (0.01, 0.05, 0.1, 1.0, 3.0), while keeping all other hyperparameters at their default values. Based on performance evaluation, the optimal configuration was determined to be n_estimators=500 and learning_rate=0.1. The fourth was the SVM classifier, which was implemented using the svm.SVC function from the scikit-learn (v1.3.2) package in Python (v3.8.19). Due to computational limitations, all hyperparameters were retained at their default values during the base algorithm selection process. The fifth was the KNN classifier, which was implemented using the neighbors.KNeighborClassifier function from the scikit-learn (v1.3.2) package in Python (v3.8.19). For this algorithm, hyperparameter tuning was performed on the n_neighbors parameter with values of 1, 3, 5, 7, and 9, while all other parameters were retained at their default settings. Based on performance evaluation, we selected n_neighbors=9 as the optimal value.

The last machine learning algorithm was the Naïve Bayes classifier, which was implemented using the naive_bayes.GaussianNB function from the scikit-learn (v1.3.2) package in Python (v3.8.19). During base algorithm selection, all parameters were kept at their default values.

The Multilayer Perceptron (MLP), which was implemented using the PyTorch package (v2.4.0) in Python (v3.8.19). The MLP consisted of three fully connected hidden layers with 1024, 512, and 64 neurons, respectively. ReLU activation was applied after each hidden layer, and dropout was used to reduce overfitting. The final output layer generated the prediction probability for virus–human PPI classification. During training, binary cross-entropy loss was used as the objective function, and the Adam optimizer was adopted for parameter optimization. The model was trained and evaluated using the same training and validation splits as the conventional machine learning algorithms.

The Feature Group Multi-Head Attention model (FeatGroup-MHAttn), which was also implemented using the PyTorch package (v2.4.0) in Python (v3.8.19). In this model, different feature categories, including ProtT5 embeddings, human protein degree, viral molecular mimicry, and PSSM embeddings, were treated as independent feature groups. Each feature group was first projected into a shared hidden dimension through a fully connected layer. The projected feature-group representations were then fed into a multi-head attention module to learn the relative contribution of different feature groups. The attention outputs were further passed through fully connected layers for final PPI classification. Similar to the MLP model, FeatGroup-MHAttn was trained using binary cross-entropy loss and optimized with the Adam optimizer under the same data-splitting strategy.

After selecting XGBoost as the base algorithm, we performed a comprehensive three-phase hyperparameter optimization. In the first phase, we systematically evaluated learning rates (0.01, 0.03, 0.05, 0.1, 0.3) and the number of estimators (ranging from 1 to 1500), identifying 0.03 and 600 as the optimal values based on loss and classification error trends (Fig. S2).

In the second phase, we conducted an exhaustive grid search of tree-specific parameters, including max_depth (3, 5, 7, 10), min_child_weight (0.1, 0.5, 1, 3, 6, 9), and gamma (0, 0.1, 0.2, 0.3, 0.4, 0.5), which yielded optimal values of 7, 0.1, and 0, respectively.

The third phase focused on subsampling strategies. We evaluated subsample (0.8, 0.9, 1.0) and colsample_bytree (0.8, 0.9, 1.0), determining that subsample=0.9 and colsample_bytree=1.0 provided the best predictive performance.

The complete set of optimized hyperparameters was provided in Table S1. Using these optimized settings, we trained vhPPIpred on the full benchmark dataset. The final model was then evaluated on independent test sets and applied to identify potential human viral receptors.

### Methods for comparison

There were nine published methods used for comparison in this study: (1) HBFormer[24], whose training dataset and published model were downloaded from https://github.com/RmQ5v/HBFormer; (2) HVPPI[21], whose training dataset and established model were obtained from http://zzdlab.com/hvppi/download.php; (3) LSTM-PHV[22], whose training dataset was obtained from http://kurata35.bio.kyutech.ac.jp/LSTM-PHV; (4) Cross-Attention_PHV[42], whose training dataset, source code and established model were obtained from https://kurata35.bio.kyutech.ac.jp/Cross-attention_PHV; (5) MultiTask-Transfer[43], whose source code was obtained from https://git.l3s.uni-hannover.de/dong/multitask-transfer; (6) TransPPI[23], whose source code was obtained from https://github.com/XiaodiYangCAU/TransPPI; (7) DeepGNHV[25], whose source code was obtained from https://github.com/bioboy0415/DeepGNHV; (8) AlphaFold-Multimer[44,45], which was run through the ColabFold implementation available at https://github.com/sokrypton/colabfold; (9) AlphaFold3[46], which was accessed through the AlphaFold Server at https://alphafoldserver.com/. All methods except LSTM-PHV and AlphaFold3, which were accessed through web servers, were executed locally on an Ubuntu 18.04.2 LTS system.

For the first six prediction methods, evaluation was conducted on samples from our benchmark dataset as well as the Yang, Zhou, and DeNovo datasets, after removing any samples that overlapped with the training samples of each respective method. For the remaining three methods, we randomly selected 50 positive and 250 negative samples from our benchmark dataset. The structures of viral and human proteins in these selected samples were predicted using AlphaFold2 with default parameters, and the resulting monomer structures were then used by DeepGNHV to predict virus-human PPIs. Separately, the complex structures of these virus-human PPIs were predicted using the AlphaFold-Multimer and AlphaFold3, also with default parameters. For each predicted complex, the interface confidence score (iPTM) was used as the predicted interaction score. For a fair comparison, vhPPIpred was retrained on the remaining samples after excluding the selected samples as well as any samples involving proteins with a sequence similarity > 0.4 to those in the selected set.

### Evaluation criteria

We introduced six common performance metrics (i.e. Accuracy, Precision, Recall, F1-Score, AUROC and AUPRC) for method evaluation and comparison. These metrics are defined as

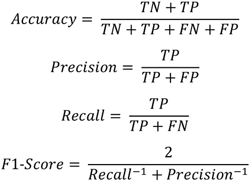

where TP, TN, FP and FN denote the number of true positives, true negatives, false positives and false negatives, respectively. All metrics were implemented using functions from the metrics module in scikit-learn (v1.3.2), including accuracy_score, precision_score, recall_score, f1_score, roc_curve, precision_reall_curve and auc.

### Permutation-based feature importance analysis

To further interpret the contribution of different feature types in vhPPIpred, we performed permutation-based feature importance analysis on our benchmark dataset. For each feature type, including ProtT5 embeddings, human protein degree, viral molecular mimicry, and PSSM embeddings, the corresponding feature values in the test set were randomly shuffled across samples, while all other features were kept unchanged. The trained vhPPIpred model was then used to predict the shuffled test set. The decreases in AUROC and AUPRC after feature permutation were calculated as the importance scores of each feature type. A larger performance decrease indicated a greater contribution of the corresponding feature type.

### Similarity between ProtT5 and PSSM embeddings

To examine whether the limited contribution of PSSM embeddings was related to information redundancy with ProtT5 embeddings, we quantified the similarity between ProtT5-derived and PSSM-derived representations in a shared latent space. This analysis was performed at both the protein level and the virus-human PPI (vhPPI) level. At the protein level, ProtT5 embeddings were standardized and then reduced from 1024 to 20 dimensions by principal component analysis (PCA) to match the dimensionality of the corresponding PSSM embeddings. Canonical correlation analysis (CCA) was then fitted between the PCA-reduced ProtT5 features and the standardized PSSM features, and both feature representations were transformed into the CCA latent space. At the protein level, virus and human proteins were analyzed separately. For each individual protein, cosine similarity was calculated between its ProtT5-derived and PSSM-derived vectors after CCA transformation. At the vhPPI level, positive and negative vhPPI pairs were analyzed separately. Pair-level ProtT5 representations consisted of concatenated PCA-reduced viral and human ProtT5 embeddings, and pair-level PSSM representations consisted of concatenated viral and human PSSM embeddings. For each vhPPI pair, cosine similarity was also calculated between its ProtT5 and PSSM embeddings in the CCA latent space.

### Sequence and motif similarity between virus and neighbor proteins

To investigate the limited contribution of viral molecular mimicry feature, we calculated the sequence similarity between the viral proteins and the neighbor proteins, defined as proteins that interact with virus-interacting human proteins. Viral proteins can interact with human proteins by mimicking their neighbor proteins[29]. Therefore, if a viral protein is known to interact with a human protein, the viral protein and the neighbor protein of this human protein were expected to exhibit sequence similarity due to this mimicking effect, and were defined as positive pairs. In contrast, if a viral protein does not interact with a human protein, the pair consisting of this viral protein and human protein’s neighbor was defined as a negative pair. Using the collected physical virus-human PPIs and the human PPI network, we randomly constructed 2,000 positive and 2,000 negative pairs. Viral proteins were queried against neighbor proteins using DIAMOND blastp, and hits with e-value smaller than 1E-6 were considered similar. The proportion of similar pairs was calculated for both positive and negative pairs.

Since virus-host interactions are often mediated by short linear motif, we further calculated motif-level similarity between viral and neighbor proteins. For each protein, 8-amino-acid motifs were generated using a sliding window with step size 1. For each virus-neighbor protein pair, sequence identity was calculated for all motif pairs, and the maximum identity was taken as the motif-level similarity. Pairs with motif-level similarity greater than 75% (6/8 amino acid identical) were considered similar. The proportion of motif-level similar pairs was calculated separately for positive and negative samples.

### Measurement of time consumption and memory usage

In this study, we systemically measured the computational performance of five prediction methods, including vhPPIpred, HBFormer, HVPPI, Cross-Attention_PHV, TransPPI and MultiTask-Transfer, in terms of memory usage and runtime. Memory usage was monitored in real time using the @profile decorator of memory-profiler package (v0.61.0) in Python (v3.8.19), enabling precise tracking of resource consumption throughout the execution process.

### Dataset for the identification of viral receptors

We downloaded 213 known interaction pairs between viral receptor binding proteins (RBPs) and human receptors from the Viral Receptor database[50]. In addition, we obtained the potential human receptorome (with RF score > 0.5) for the human-infecting virome from Zhang’s dataset[51], accessed on September 4, 2021.

### Annotations of predicted viral receptor candidates

To characterize the predicted viral receptor candidates, we selected the top 10 predictions for seven viruses whose known receptors were ranked within the top 10 by vhPPIpred. A total of 19 human cell membrane proteins were included among these selected predictions. Annotations, including gene symbols, protein functions, subcellular locations, and Gene Ontology (GO) terms, were manually retrieved from the UniProt database (accessed on May 14, 2026). Based on predefined keywords in the functional annotations, the proteins were classified into six functional categories: (1) known viral receptors, (2) virus-associated host factors, (3) viral entry/trafficking, (4) cell adhesion/junction, (5) surface receptor signaling, and (6) others.

### Visualization of results

All figures in this study were generated using Python (3.8.19) in combination with Pandas (v2.0.3), NumPy (v1.24.4) and Matplotlib (v3.7.5). Network visualization was performed using Cytoscape (v3.10.2).

## Acknowledgments

**General:** We thank members in PengLab for helpful discussions on the manuscript.

## Author contributions

Conceptualization: Z.Z., and Y.P. Methodology: Z.Z. Investigation: Z.Z., and Y.F. Visualization: Z.Z., and Y.F. Supervision: Z.Z., X.M., and Y.P. Writing—original draft: Z.Z., and Y.P. Writing—review & editing: Z.Z., X.M., and Y.P.

## Data availability

All data used in the study are available in supplementary materials. The vhPPIpred is publicly available at https://github.com/ZyuanZhang/vhPPIpred.

