## Supplementary material for "Improved prediction of virus-human protein-protein interactions by incorporating network topology and viral molecular mimicry": Fig S1-S4 and Table S1-S5

Zhiyuan Zhang *et al.*

**This file includes:**

Figs. S1 to S4
Tables S1 to S5
References

**Fig. S1. Pairwise Jaccard indices of human protein clusters across the six groups in the benchmark dataset.** Jaccard indices[1] represent the overlap between any two groups of benchmark dataset.

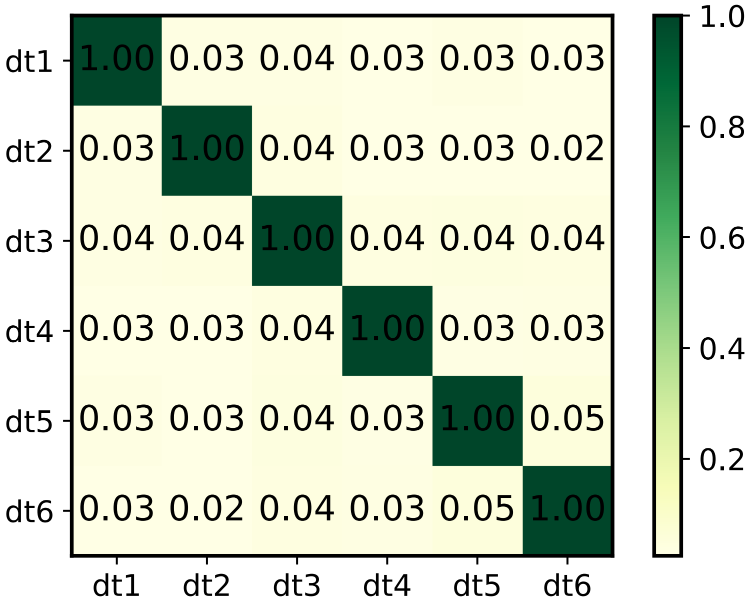

**Fig. S2. Training and validation log loss and classification error across n_estimators at different learning rates in vhPPIpred.** We tracked the training and validation log loss **(A)** and classification error **(B)** using the evals_result function of XGBoost (v2.1.1)[2] in Python (v3.8.19)[3] For each learning rate (LR), results were averaged across folds in six rounds of 5-fold cross-validation to generate the final curves.

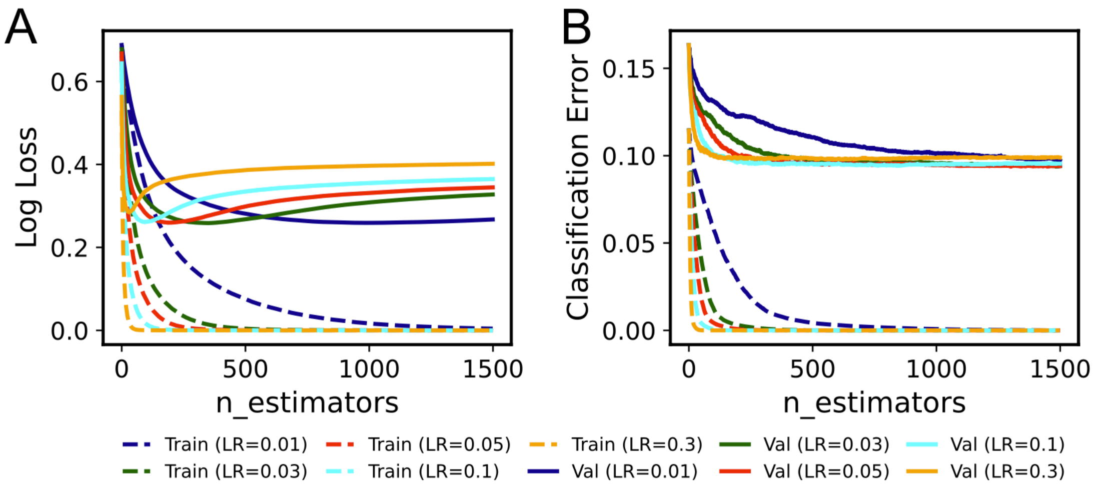

**Fig. S3. Prediction scores of AlphaFold2, AlphaFold-Multimer and AlphaFold3 on 300 randomly selected virus-human PPIs. (A)** The pLDDT distribution of predicted structures of human and virus monomer proteins predicted by AlphaFold2[4]. The iPTM distribution of predicted dimer structures of virus-human PPIs predicted by AlphaFold-Multimer[5] **(B)** and AlphaFold3[6]**(C)**.

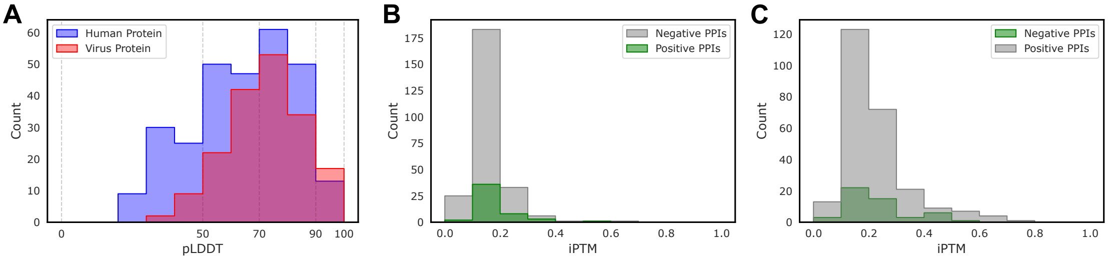

**Fig. S4. Pipeline for negative sample construction, applied to both Zhou's dataset and the DeNovo dataset.**

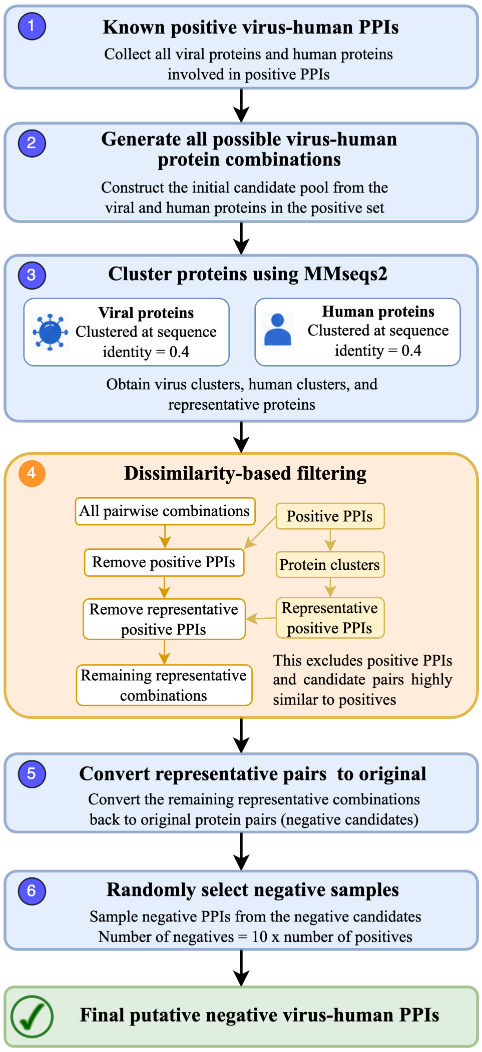

**Table S1. Optimal hyperparameter combinations of XGBoost.** Key hyperparameters of XGBoost and their candidate values are listed. The optimal combination was determined through grid search with 5-fold cross-validation on training datasets of benchmark dataset, based on validation performance.

| **Parameter** | **Value Range** | **Optimal Value** |
|---|---|---|
| learning_rate | [0.01, 0.03, 0.05, 0.1, 0.3] | 0.03 |
| n_estimators | 1-1500 | 600 |
| max_depth | [3, 5, 7, 10] | 7 |
| min_child_weight | [0.1, 0.5, 1, 3, 6, 9] | 0.1 |
| gamma | [0, 0.1,0.2,0.3,0.4,0.5] | 0 |
| subsample | [0.8, 0.9, 1.0] | 0.9 |
| colsample_bytree | [0.8, 0.9, 1.0] | 1.0 |

**Table S2. Comparison of prediction methods on benchmark dataset.** Methods marked with “#” were evaluated on benchmark dataset after removing overlapping PPIs with the training set of the corresponding established models. Methods marked with “*” indicated models that were retrained and evaluated on benchmark dataset.

| **Methods** | **Accuracy** | **Precision** | **Recall** | **F1-Score** | **AUROC** | **AUPRC** |
|---|---|---|---|---|---|---|
| vhPPIpred | 0.913 | **0.952** | 0.050 | 0.094 | **0.921** | **0.680** |
| HBFormer[7] | **0.916** | 0.768 | 0.106 | 0.186 | 0.774 | 0.372 |
| HVPPI[8] | 0.915 | 0.754 | 0.098 | 0.173 | 0.854 | 0.515 |
| LSTM-PHV[9] | 0.797 | 0.262 | **0.678** | **0.378** | 0.699 | 0.193 |
| Cross-Attention_PHV[10] | 0.848 | 0.149 | 0.143 | 0.146 | 0.510 | 0.109 |
| HBFormer # | 0.949 | 0.670 | 0.070 | 0.126 | 0.750 | 0.251 |
| HVPPI # | **0.965** | 0.315 | 0.043 | 0.075 | 0.793 | 0.216 |
| LSTM-PHV # | 0.795 | 0.137 | **0.540** | **0.219** | 0.597 | 0.107 |
| Cross-Attention_PHV # | 0.856 | 0.119 | 0.128 | 0.123 | 0.492 | 0.088 |
| MultiTask-Transfer *[11] | 0.886 | 0.372 | 0.376 | 0.374 | 0.693 | 0.376 |
| TransPPI *[12] | 0.912 | 0.527 | **0.410** | **0.458** | 0.809 | 0.446 |
| Cross-Attention_PHV * | 0.872 | 0.318 | 0.293 | 0.289 | 0.759 | 0.286 |

**Table S3. Performance comparison of vhPPIpred with structure-based virus-human PPI prediction methods on a subset of benchmark dataset (50 positive and 250 negative PPIs).** Accuracy, Precision, Recall, and F1-Score in vhPPIpred and DeepGNHV[13] were computed using a threshold of 0.5, while the corresponding values in AlphaFold-Multimer and AlphaFold3 were based the threshold yielding the highest Accuracy. Additionally, vhPPIpred was retrained on remaining samples of benchmark dataset that removed proteins similar to the selected samples.

| | **Accuray** | **Precision** | **Recall** | **F1-Score** | **AUROC** | **AUPRC** |
|---|---|---|---|---|---|---|
| vhPPIpred | **0.933** | **0.813** | **0.780** | **0.796** | **0.972** | **0.854** |
| AlphaFold-Multimer | 0.833 | 0.500 | 0.060 | 0.107 | 0.535 | 0.213 |
| AlphaFold3 | 0.797 | 0.077 | 0.020 | 0.032 | 0.502 | 0.180 |
| DeepGNHV | 0.797 | 0.415 | 0.540 | 0.470 | 0.692 | 0.564 |

**Table S4. Performance comparison of vhPPIpred integrating protein embeddings generated by different pretrained protein language models (PLM).**

| PLM | Accuray | Precision | Recall | F1-Score | AUROC | AUPRC |
|---|---|---|---|---|---|---|
| ProtT5[14] | **0.905** | **0.491** | **0.732** | **0.585** | **0.919** | **0.670** |
| ProtBert[14] | 0.874 | 0.400 | 0.728 | 0.515 | 0.893 | 0.593 |
| ESM-2[15] | 0.893 | 0.453 | 0.731 | 0.556 | 0.907 | 0.622 |

**Table S5. Keywords used to filter physical interactions across human protein-protein interaction databases.** This table summarizes the specific keywords employed to identify and retain only physical interactions from various human PPI databases.

| **Database** | **Keywords** | **Download date** | **Version** |
|---|---|---|---|
| Mentha[16] | | April 25, 2022 | v1.0 |
| MINT[17] | "direct interaction", "disulfide bond", "association", "covalent binding" | May 5, 2022 | v1.0 |
| IntAct[18] | "direct interaction", "self interaction", "association", "covalent binding" | May 5, 2022 | v4.2.3.2 |
| STRING[19] | 9606.protein.physical.links.v11.5.txt.gz | May 16, 2022 | v11.5 |
| BioPlex[20] | | May 30, 2022 | v3.0 |
| BioGRID[21] | "direct interaction" | June 30, 2022 | v4.4.208 |

21. Oughtred R, Rust J, Chang C, Breitkreutz B, Stark C, Willems A, et al. The

BioGRID database: A comprehensive biomedical resource of curated protein, genetic, and chemical interactions. Protein Sci. 2021;30:187–200. https://doi.org/10.1002/pro.3978
